# Gallein and isoniazid act synergistically to attenuate *Mycobacterium tuberculosis* growth in human macrophages

**DOI:** 10.1101/2024.01.10.574965

**Authors:** Ramesh Rijal, Richard H. Gomer

**Author notes:** Corresponding author (RR) or (RHG).

## Abstract

*Mycobacterium tuberculosis* (*Mtb*), the bacterium that causes tuberculosis (TB), can be difficult to treat because of drug resistance. Increased intracellular polyphosphate (polyP) in *Mtb* enhances resistance to antibiotics, and capsular polyP in *Neisseria gonorrhoeae* potentiates resistance to antimicrobials. The mechanism by which bacteria utilize polyP to adapt to antimicrobial pressure is not known. In this study, we found that *Mtb* adapts to the TB frontline antibiotic isoniazid (INH) by enhancing the accumulation of cellular, extracellular, and cell surface polyP. Gallein, a broad-spectrum inhibitor of the polyphosphate kinase that synthesizes polyP, prevents this INH-induced increase in extracellular and cell surface polyP levels. Gallein and INH work synergistically to attenuate *Mtb*’s ability to grow in *in vitro* culture and within human macrophages. *Mtb* when exposed to INH, and in the presence of INH, gallein inhibits cell envelope formation in most but not all *Mtb* cells. Metabolomics indicated that INH or gallein have a modest impact on levels of *Mtb* metabolites, but when used in combination, they significantly reduce levels of metabolites involved in cell envelope synthesis and amino acid, carbohydrate, and nucleoside metabolism, revealing a synergistic effect. These data suggest that gallein represents a promising avenue to potentiate the treatment of TB.

**Author summary:** *Mycobacterium tuberculosis* (*Mtb*) is the causative agent of tuberculosis (TB), which is responsible for more deaths than any other infectious disease. The alarming prevalence of drug-resistant *Mtb* strains has further exacerbated this global health crisis. Some pathogenic bacteria such as *Mtb* appear to increase levels of polyphosphate as a defense against antibiotics. We found that gallein, a small molecule inhibitor of bacterial polyphosphate kinases, strongly potentiates the ability of the frontline anti-tuberculosis drug isoniazid to inhibit the growth of *Mtb* both alone and in human macrophages. This has unveiled vulnerabilities in *Mtb* that could be strategically leveraged to reverse INH resistance.

## Introduction

Tuberculosis (TB) remains a significant global public health challenge, and in 2022, the causative bacterium *Mycobacterium tuberculosis* (*Mtb*) was responsible for approximately 1.6 million deaths worldwide [1]. During infection, *Mtb* encounters a variety of stressors originating from the host, and in response, employs adaptive physiological mechanisms to endure these stresses, promoting both resistance to antibiotics and the development of drug resistance [2-6]. These complexities not only demand prolonged treatment regimens but also contribute to the emergence of drug-resistant *Mtb* strains [1]. Notably, resistance to the primary antibiotic, isoniazid (INH), is a prevalent form of monoresistance in *Mtb*, which is associated with treatment failures and the emergence of multidrug-resistant TB [1]. *Mtb*’s resistance to the majority of antibiotics is attributed to the thickening of the cell envelope [7, 8], the activation of enzymes that modify antibiotics or their targets, and the action of efflux pumps [9, 10]. These mechanisms collectively reduce the efficacy of antibiotics.

Polyphosphate (polyP) is a chain of phosphate residues and is present in all kingdoms of life [11]. PolyP metabolism has been linked to the virulence of pathogens such as *Mtb, Burkholderia mallei*, *Pseudomonas aeruginosa*, *Salmonella enterica*, and *Shigella flexneri* [12- 15]. A highly conserved bacterial enzyme, polyphosphate kinase (PPK), synthesizes polyP from ATP, while polyP levels are regulated by the action of exopolyphosphatase (PPX), an enzyme that removes terminal phosphate residues from a polyP chain [16]. The *Mtb* genome encodes two PPKs, PPK1 (Rv2984) and PPK2 (Rv3232c), as well as two PPXs, PPX1 (Rv0496) and PPX2 (Rv1026) [17].

Pathogenic bacteria lacking PPK or having reduced PPK levels exhibit defects in stress response, quorum sensing, growth, survival, and virulence [13, 18-24]. For instance, intracellular polyP is necessary for the survival of *Mtb* in host cells [14, 22, 25, 26], and deletion of PPK1 in *M. smegmatis* attenuates the survival of ingested *M. smegmatis* in human macrophages [27].

Conversely, increased intracellular polyP in *Mtb* causes increased resistance to antibiotics [14, 26, 28-30]. In addition to intracellular polyP, bacteria also have extracellular polyP. The pathogenic bacterium *Neisseria gonorrhoeae* has polyP in its capsule, and the polyP potentiates resistance to antimicrobials [23, 24]. We observed that both *Mtb* and *M. smegmatis* accumulate extracellular polyP [27]. Treatment of *Mtb*-infected macrophages with a polyP-degrading recombinant exopolyphosphatase (ScPPX) reduced the *Mtb* burden in macrophages, suggesting that both intracellular and extracellular polyP potentiate the survival of *Mtb* in host cells [27].

Given the absence of PPK enzymes in humans [31], bacterial PPKs could serve as potential targets for antituberculosis therapeutics. The small molecule gallein [32] is a broad-spectrum PPK inhibitor [33, 34], and in this report, we find that gallein strongly potentiates the ability of INH to inhibit *Mtb* growth alone and in human macrophages.

## Results

### Gallein enhances the INH-mediated inhibition of *Mtb* growth

PPK1 and PPK2, enzymes which synthesize polyP, are both necessary for *Mtb* viability [21, 35]. Ellagic acid derivatives from a medicinal plant inhibit PPK1 in *Pseudomonas aeruginosa* [36]. Gallein, a small molecule with similarity to ellagic acid, was identified as a potent inhibitor of both PPK1 and PPK2 in *P. aeruginosa* [33]. Gallein also inhibits Gβγ subunit signaling in mammalian cells [37, 38]. To determine if gallein affects *Mtb*, cells were cultured in the presence of different concentrations of gallein. Gallein at 0.005 µM, 0.05 µM, or 0.5 µM did not significantly affect *Mtb* growth, as assessed by an increase in OD_600_ values (Figures S1 A-C), while 5 µM gallein slightly slowed growth, and 50 µM gallein inhibited growth by approximately 80% (Figures 1 A and B). In the absence of gallein, 1 µg/ml INH caused a partial but not complete reduction in *Mtb* growth (Figure S1 D), and this effect was potentiated by 5 and 50 µM gallein (Figures 1 A and B). The addition of 0.005 µM, 0.05 µM, or 0.5 µM gallein did not significantly enhance the effect of 1 µg/ml INH (Figures S1 A-C). To determine if the reduced growth with INH and/or 5 and 50 µM gallein causes a permanent effect on *Mtb*, the day 14 cultures were washed and resuspended in medium without INH or gallein and the growth was monitored (Figure S1 E). Surprisingly, exposure to INH caused *Mtb* cells to grow faster and exposure to gallein caused a slight reduction in growth. Together, these data indicate that a 14 day exposure of *Mtb* to 1 µg/ml INH and 5 or 50 µM gallein is bacteriostatic but not bactericidal.

**Figure 1:**
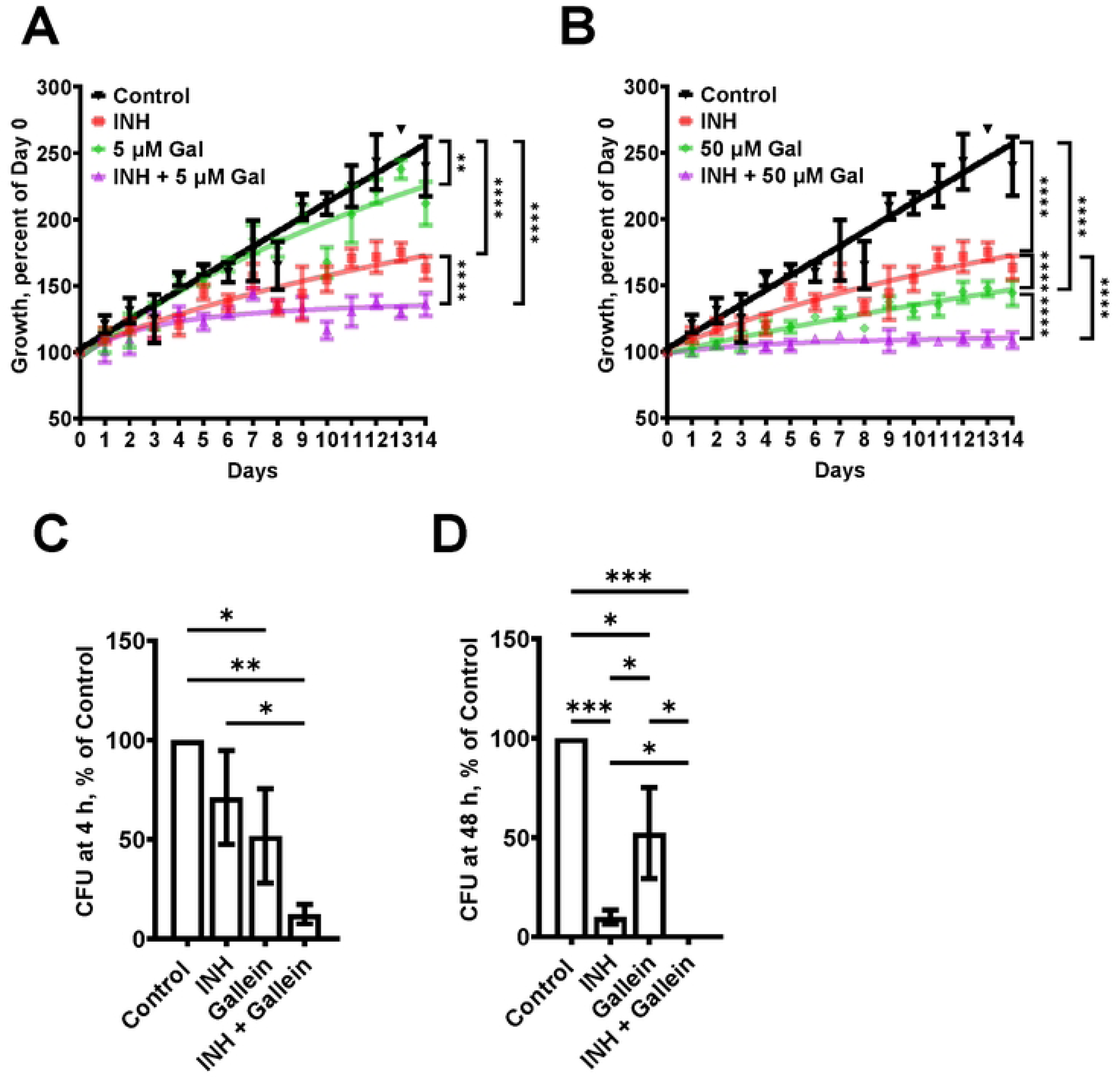
Gallein potentiates the ability of INH to inhibit *Mtb* growth both in *in vitro* culture and within macrophages. (A and B) *Mtb* cultures were grown for 14 days in the absence (Control) or presence of 1 µg/ml isoniazid (INH) and/or 5 µM (A) or 50 µM (B) gallein. The OD_600_ was measured daily, and growth was determined as a percentage of Day 0 OD_600_. (C and D) Viable ingested *Mtb* in macrophages, in the absence (Control) or presence of 1 µg/ml INH and/or 5 µM gallein, was determined as colony-forming units (CFU) at 4 hours (C) and 48 hours (D) after ingestion. CFU in the control was considered 100%. All values are mean ± SEM of three (A and B) and four (2 females and 2 males) (C and D) independent experiments. * P < 0.05; ** P < 0.01; *** P < 0.001, **** P < 0.0001 (Two-way ANOVA with Dunnett’s multiple comparisons test for A and B, and Mann-Whitney test for C and D).

In patients with tuberculosis, *Mtb* bacteria can be ingested by macrophages, and are able to survive inside the macrophages [39]. To determine if gallein affects the ability of *Mtb* to survive in macrophages, human macrophages, derived from circulating monocytes from healthy donors, were incubated with *Mtb* for 2 hours, and the non-ingested *Mtb* were removed. The macrophages were then incubated with gallein and/or INH, and at 4 and 48 hours after adding *Mtb* the macrophages were lysed with a detergent (0.1% Triton X-100) that does not kill *Mtb*, and the bacteria were plated and colonies were counted. At 48 but not 4 hours, INH decreased the viability of ingested *Mtb* (Figures 1 C and D). At both times, gallein decreased ingested *Mtb* viability compared to control, and in the presence of INH significantly decreased the viability of ingested *Mtb*, with no detected surviving bacteria at 48 hours (Figures 1 C and D). Together, these results suggest that for both free *Mtb* and *Mtb* in macrophages, gallein inhibits growth and enhances INH’s ability to inhibit growth.

### Isoniazid increases the accumulation of polyP

Infection of host cells with *Mtb* leads to a resistance of the *Mtb* to INH [4, 40, 41], and increased intracellular polyP in *Mtb* causes increased resistance to antibiotics [14, 26, 28-30]. To determine if exposure of *Mtb* to INH potentiates accumulation of polyP (28, 29), *Mtb* cells were cultured in the presence of INH. After exposure to INH for 21 days, the *Mtb* cells were fixed without permeabilizing the cells and then stained with the GFP-tagged polyP binding domain of *Escherichia coli* PPX (GFP-PPX) [42]. For unknown reasons, the cells showed a wide range of staining intensities (Figures 2 A and B). INH concentrations from 0.1 to 100 µg/ml increased the average amount of cell-surface polyP (Figures 2 A and B). A 1 day exposure of *Mtb* to 1 µg/ml INH also increased cell-surface and total cellular polyP (Figures 2 C - E). We previously observed that *Mtb* cells accumulate extracellular polyP [27], and at 1 day INH also increased the accumulation of extracellular polyP (Figure 2 F). At 5 and 14 days, 1 µg/ml INH significantly increased cell-surface, total cellular, and extracellular polyP (Figures 2 G - L). At 14 days, *Mtb* controls had, using an arbitrary cutoff, 23 % of cells with low levels of cell surface polyP, cells treated with gallein in the presence or absence of INH had 43% of cells with low levels of cell surface polyP, but none of the cells treated with INH had low levels of cell surface polyP (Figure S2A). Together, these findings suggest that INH induces the accumulation of cellular, extracellular, and cell surface polyP.

**Figure 2:**
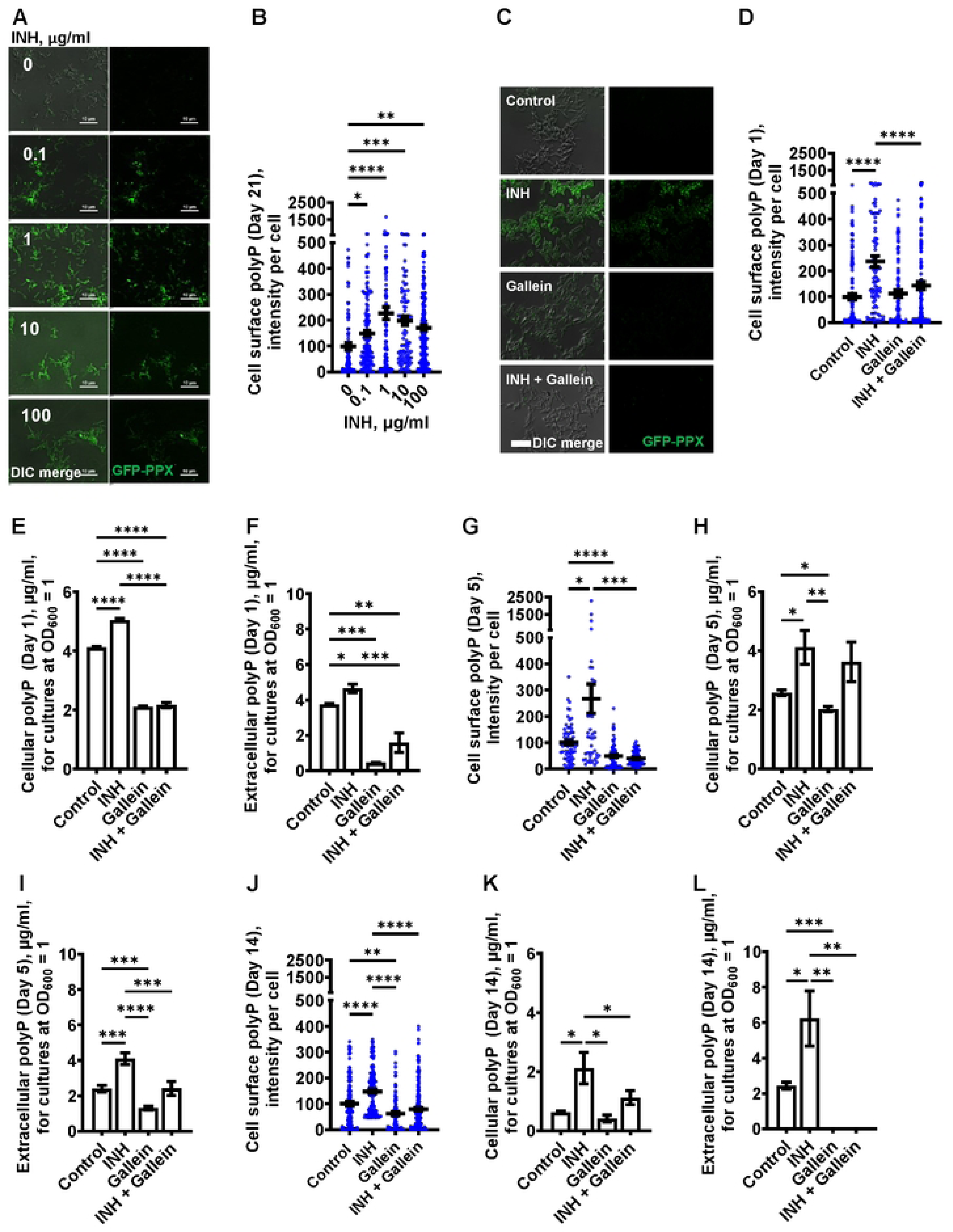
INH induces accumulation of *Mtb* cell surface, cellular, and extracellular polyP. (A) *Mtb* were cultured for 21 days in the presence of the indicated concentrations of INH and then stained with GFP-PPX. Differential interference contrast (DIC) merged with fluorescence are at the left, and fluorescence images are at the right. Representative images from at least three independent experiments are shown. Bars are 10 µm. (B) GFP-PPX staining intensity per cell, as a measure of *Mtb* cell surface polyP per cell, was determined from panel A. Black bars are means. In B, D, G, and J, the staining intensities are in arbitrary units with the control averages set to 100. (C) DIC merge and fluorescence images of *Mtb* grown for 1 day in the absence or presence of 1 µg/ml INH and/or 5 µM gallein and stained with GFP-PPX are displayed. Representative images from at least three independent experiments are shown. Bar is10 µm. (D) GFP-PPX staining intensity per cell, as a measure of *Mtb* cell surface polyP per cell, was determined from panel C. Black bars are means. (E) *Mtb* were grown for 1 day in the absence or presence of 1 µg/ml INH and/or 5 µM gallein. The OD_600_ was measured, cells were lysed, and cellular polyP levels for cultures at OD_600_=1 were determined. (F) *Mtb* were grown for 1 day in the absence or presence of 1 µg/ml INH and/or 5 µM gallein. The OD_600_ was measured, and supernatant conditioned medium was collected. Extracellular polyP levels for cultures at OD_600_=1 were determined. (G - L) *Mtb* were grown for 5 days (G – I) or 14 days (J – L) in the absence or presence of 1 µg/ml INH and/or 5 µM gallein. Levels of cell surface polyP (G and J), cellular polyP (H and K), and extracellular polyP (I and L) were determined as described in (C – F). All values are mean ± SEM of three independent experiments. * P < 0.05; ** P < 0.01; *** P < 0.001, **** P < 0.0001 (One-way ANOVA with Tukey’s multiple comparisons test).

### Gallein prevents the INH-induced accumulation of extracellular and cell surface polyP

One possible mechanism for the effect of gallein on *Mtb* growth is that gallein, by inhibiting PPKs, decreases polyP levels. At 1 day, in the absence of INH, gallein did not decrease cell surface polyP, but did block the INH-induced increase in cell surface polyP, bringing the cell surface polyP levels to levels comparable to control cells (Figure 2 D). Gallein decreased cellular polyP in both the absence and presence of INH (Figure 2 E). Gallein also decreased extracellular polyP in both the absence and presence of INH (Figure 2 F). INH and gallein did not significantly affect levels of *Mtb ppk1* and *ppk2* mRNAs (Figure S3), suggesting that the effects on polyP levels are not mediated by changes in the levels of these mRNAs. At 5 days, gallein significantly decreased cell surface polyP in both the absence or in the presence of INH (Figure 2 G). At 5 days, gallein decreased cellular polyP in the absence of INH (Figure 2 H). Gallein also decreased extracellular polyP in both the absence and presence of INH (Figure 2 I). In the presence of INH, gallein decreased extracellular polyP to levels comparable to control cells (Figure 2 I). At 14 days, gallein significantly decreased cell surface polyP in both the absence or in the presence of INH (Figure 2 J). At 14 days, gallein did not significantly decrease cellular polyP in the absence of INH (Figure 2 K), but significantly decreased cellular polyP in the presence of INH (Figure 2 K). Gallein decreased extracellular polyP in both the absence and presence of INH (Figure 2 L). At 14 days, in the absence or presence of INH, gallein decreased extracellular polyP to an undetectable level (Figure 2 L). At 14 days, in the presence of the combination of INH and gallein, cellular debris was visible, indicating cell death (Figure S2 B). For unknown reasons, in control cells, cellular and extracellular polyP levels tended to decrease with the age of the cultures. Together, these results suggest that INH increases total, cell surface, and extracellular polyP, and that gallein blocks this effect except for total polyP at day 5.

### Gallein prevents INH-induced thickening of the *Mtb* cell envelope

When exposed to INH, *Mtb* undergoes cell envelope thickening and reduces cell envelope permeability as an adaptive response to withstand INH [28]. To investigate whether gallein can reverse the effects of INH on cell envelope morphology, *Mtb* was treated with 1 µg/ml INH and/or 5 µM gallein for 14 days, and the *Mtb* cell wall was visualized using transmission electron microscopy (TEM). As previously observed [28], INH increased *Mtb* cell envelope thickness (Figures 3 A and B). Gallein alone did not significantly affect cell envelope thickness. For cells exposed to both INH and gallein, the average cell envelope thickness was comparable to control cells, but there were two distinct populations of *Mtb* cells (Figures 3 A and B). For 43.0 ± 2.6% (mean ± SEM, n=3) of the *Mtb* in the presence of INH and gallein, there was a detectable cell envelope, while the remaining *Mtb* had no detectable cell envelope (Figures 3 A and B). These results suggest that in the absence of INH, gallein does not inhibit growth by affecting cell envelope thickness, and that the combination of INH and gallein causes the formation of two populations of cells, one with a detectable cell envelope (which for many of these cells is thicker than that of control cells), and one with a compromised cell envelope.

**Figure 3:**
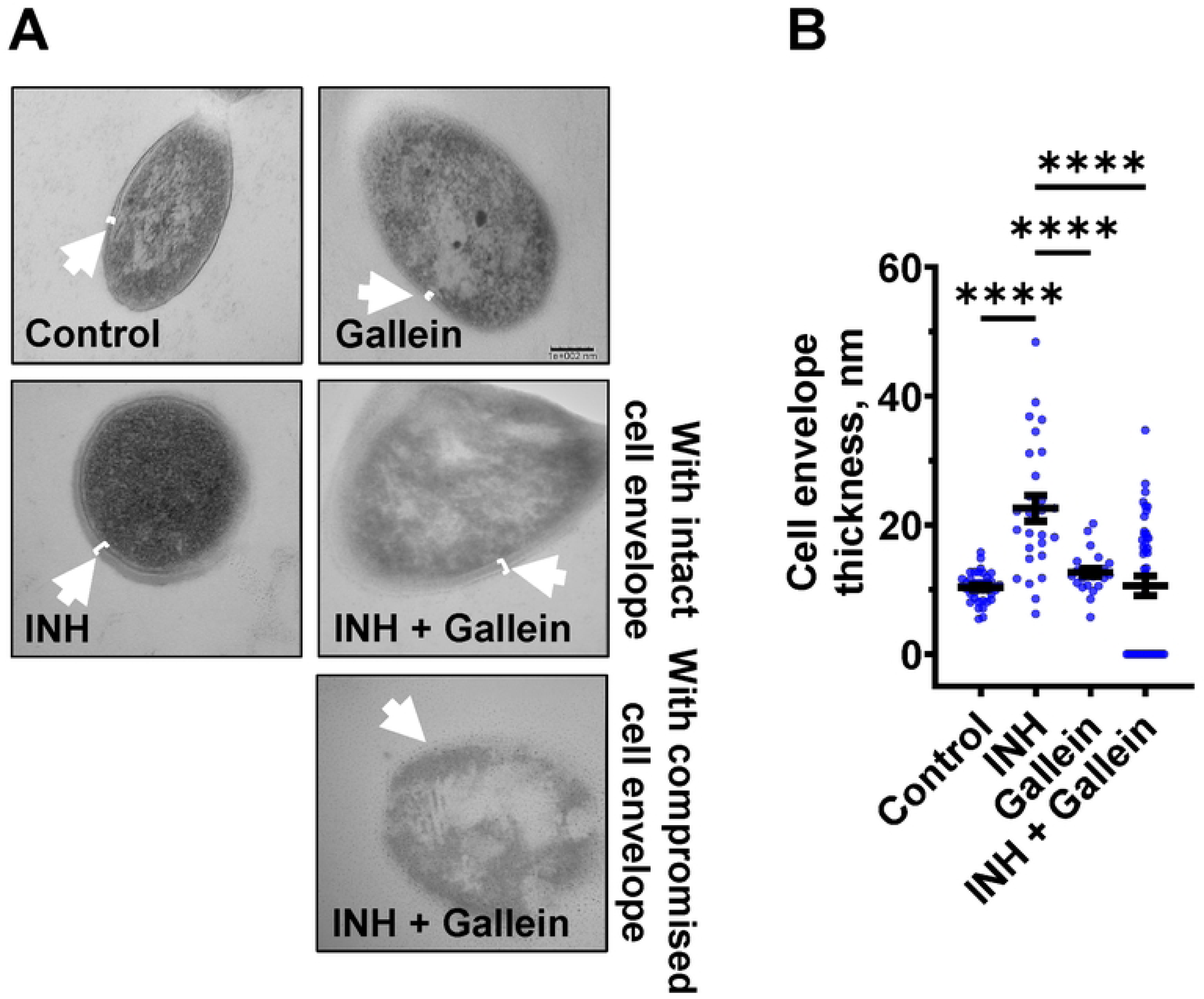
Gallein inhibits INH-induced *Mtb* cell envelope thickening. (A) Transmission electron microscopy images of *Mtb* treated without or with 1 µg/ml isoniazid (INH) and/or 5 µM gallein for 14 days. Representative images are from at least three independent experiments. The arrows indicate the cell envelope. Bar is 100 nm. (B) Quantification of cell envelope thickness. Values are mean ± SEM of at least 25 cells from three independent experiments. **** P < 0.0001 (One-way ANOVA with Tukey’s multiple comparisons test).

### Gallein and INH work synergistically to downregulate key metabolic pathways

To evaluate the impact of gallein on *Mtb* metabolism, *Mtb* were exposed to 1 µg/ml INH and/or 5 µM gallein for 24 hours, then incubated with a viability dye for 12 hours, and the resulting fluorescence signal was measured. The conversion of the non-fluorescent dye resazurin to the fluorescent resorufin product serves as a measure of metabolism [43, 44], and is also used to assess the susceptibility of *Mtb* to antimicrobial compounds [45]. In comparison to the control, *Mtb* exposed to INH showed increased metabolic activity (Figure S4A). Gallein decreased metabolic activity in the presence or absence of INH (Figure S4A).

Three clinically used tuberculosis drugs, namely INH, rifampicin (RIF), and streptomycin (STREP), induce a common pattern of metabolic alterations [46]. To determine how gallein affects metabolites, *Mtb* was exposed to 1 µg/ml INH and/or 5 µM gallein for 24 hours. *Mtb* metabolites were extracted and subjected to untargeted metabolomics analysis, which identified a total of 119 metabolites. Partial least-squares discriminant analysis (PLS-DA) revealed a clear distinction between the *Mtb* populations treated with gallein and INH in comparison to the control. Component 2, responsible for the largest proportion of the total variance in metabolites (8.6%), placed the gallein and INH treated samples considerably apart from the control, gallein, or INH treated samples (Figure S4B). This observation suggests that the metabolic changes induced by gallein and INH were more substantial than those caused by gallein or INH alone (Figure S4B). A heatmap also indicated that although INH alone and gallein alone have some effect on metabolites, the combination of gallein and INH has a more profound effect (Figure 4A).

**Figure 4:**
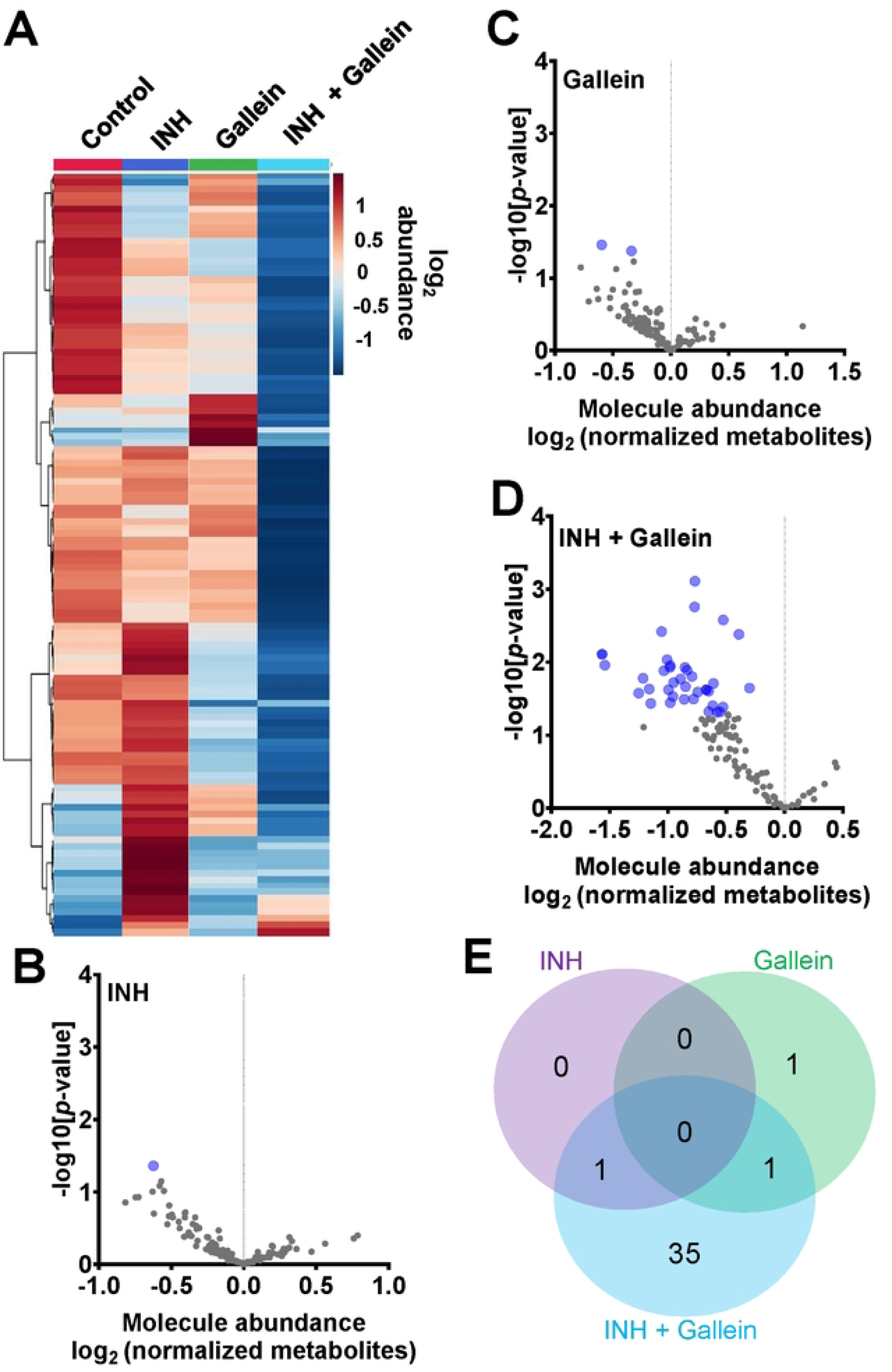
INH and gallein work synergistically to reduce metabolite levels. (A) Hierarchical clustering heatmap of 119 metabolites from *Mtb* treated with or without 1 µg/ml INH and/or 5 µM gallein for 24 hours. Rows represent the mean abundance of individual metabolites detected in all three independent experiments, normalized to total protein content. The data were generated using the metabolomics data analysis tool MetaboAnalyst (https://www.metaboanalyst.ca/), and values are depicted on a log2 scale. (B - D) The abundance of individual metabolites detected in all three independent experiments was normalized to total protein content, and the mean fold change of normalized abundance of individual metabolites relative to the control on a log2 scale was plotted against the −log10 [p-value] to generate volcano plots. Blue dots indicate metabolites with relative levels having P < 0.05 (Student’s t-test). (E) A Venn diagram depicts unique and common metabolites that were significantly reduced in (B – D). All values represent the mean ± SEM of three independent experiments.

As previously observed using a variety of INH concentrations [47], 1 µg/ ml INH significantly reduced the levels of nicotinamide adenine dinucleotide (NAD^+^) (Figure 4B and Table 1). Other workers also found that 6.4 µg/ml INH alters levels of many metabolites [46]. We observed that gallein significantly reduced the levels of deoxythymidine diphosphate (dTDP) and biotin (Figure 4C and Table 1). The combination of gallein and INH significantly reduced the levels of 37 metabolites (Figure 4D and Table 1) including NAD^+^ and dTDP but not biotin (Figure 4E). These pathways are detailed in Table 2 and include nucleoside and nucleotide biosynthesis and degradation, carrier, cofactor, and vitamin biosynthesis, amino acid biosynthesis and degradation, carbohydrate biosynthesis and degradation, aminoacyl tRNA charging, cell structure biosynthesis, metabolic regulator biosynthesis, fermentation, and fatty acid and lipid degradation pathways. Gallein, INH, or the combination of gallein and INH, did not significantly upregulate any detectable metabolite. These data indicate that gallein and INH work synergistically to impact the majority of key metabolic pathways, contributing to the inhibition of *Mtb* growth.

**Table 1.**
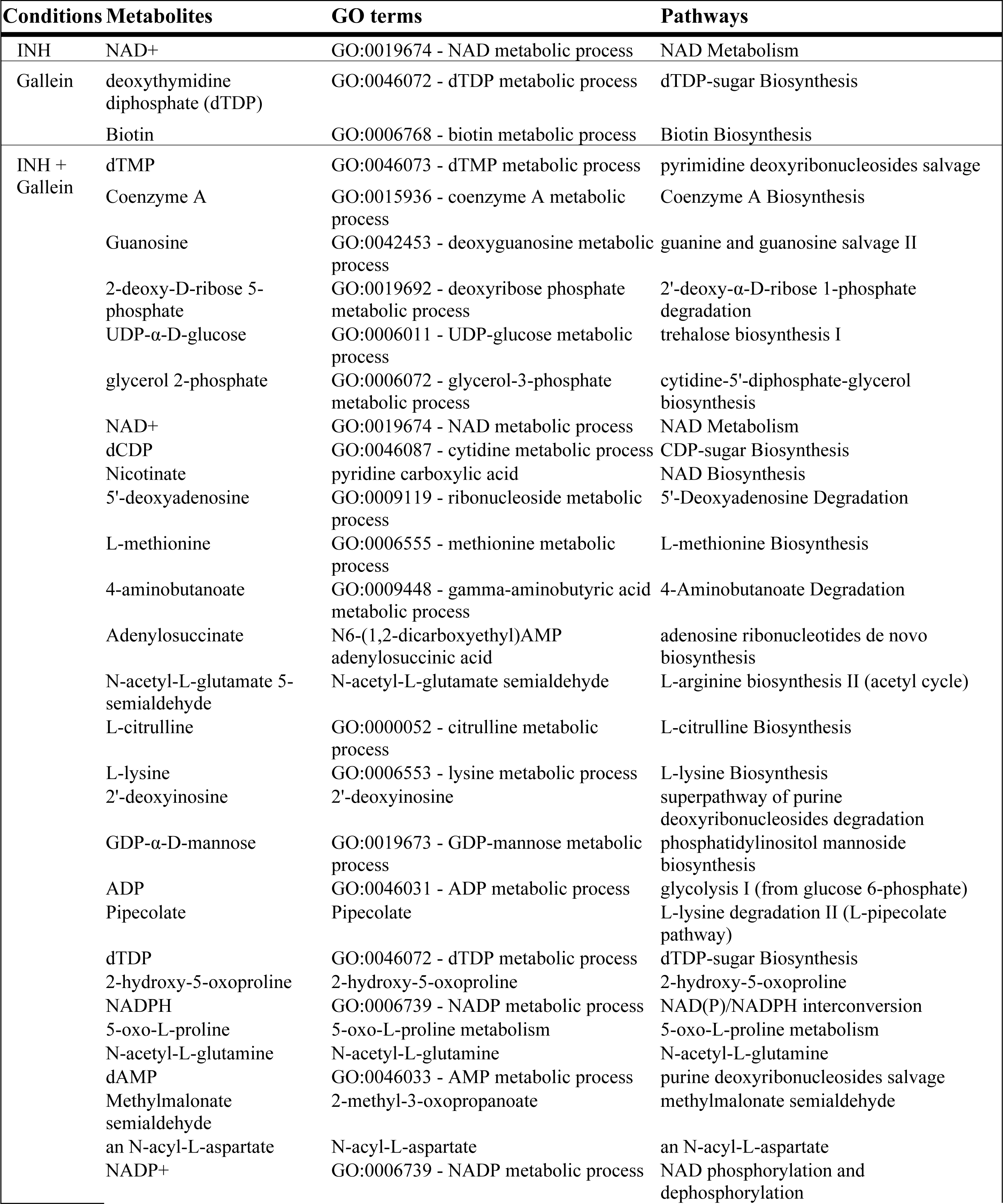
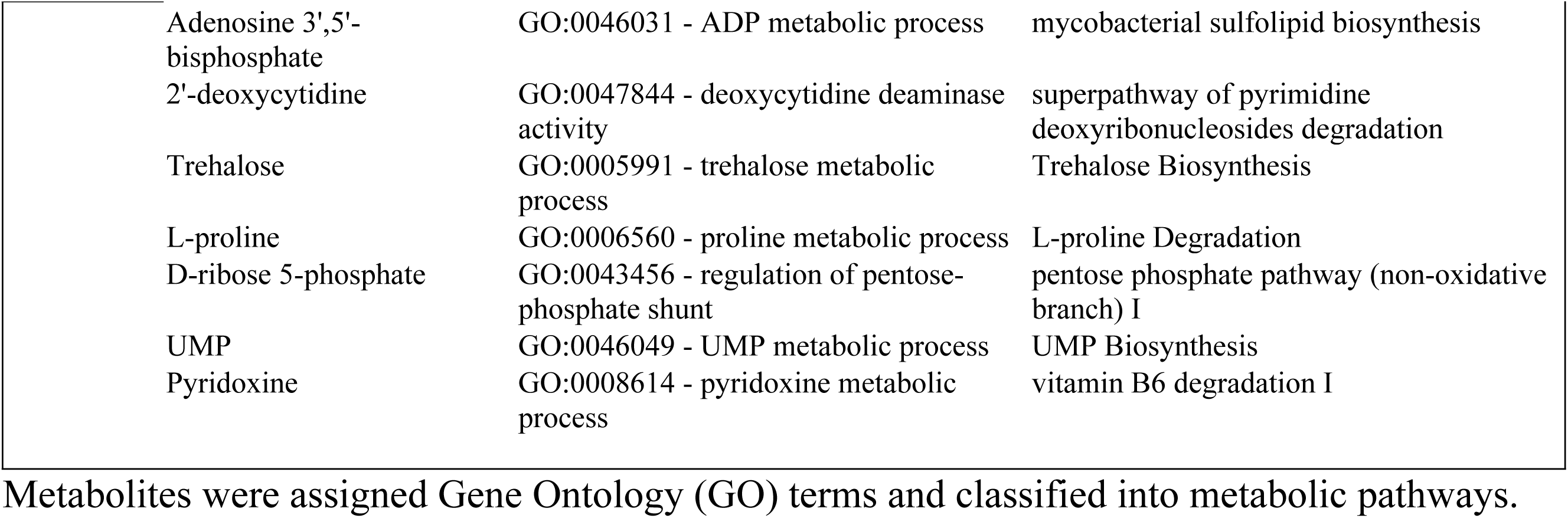
Metabolites significantly altered in *Mtb* treated with INH and/or gallein, as compared to the control.

**Table 2.** Characterization of metabolites significantly altered in *Mtb* treated with the combination of INH and gallein compared to control.

| Metabolites | Pathways |
| --- | --- |
| UMP<br>dCDP<br>dTMP<br>dTDP<br>Guanosine<br>Adenosine<br>Adenylosuccinate<br>ADP | Nucleoside and Nucleotide Biosynthesis |
| NADP<sup>+</sup><br>NAD<sup>+</sup><br>NADPH<br>L-methionine<br>Nicotinate | Carrier, Cofactor, and Vitamin Biosynthesis |
| L-proline<br>L-citrulline<br>L-methionine<br>L-lysine<br>N-acetyl-L-glutamate 5-seminaldehyde | Amino acid Biosynthesis |
| UDP-α-D-glucose<br>GDP-α-D-glucose<br>D-ribose 5-phosphate<br>Trehalose | Carbohydrate Biosynthesis |
| Methionine<br>L-lysine<br>L-proline | Aminoacyl tRNA charging |
| D-ribose 5-phosphate<br>UDP-α-D-glucose | Cell Structure Biosynthesis |
| UDP-α-D-glucose<br>Trehalose | Metabolic Regulator Biosynthesis |
| 5-oxo-L-proline<br>D-ribose 5-phosphate | Fermentation |
| 2-deoxy-D-ribose 5-phosphate<br>Adenosine<br>Guanosine<br>D-ribose 5-phosphate | Nucleoside and Nucleotide Degradation |
| UDP-α-D-glucose<br>Trehalose<br>Glycerol | Carbohydrate Degradation |
| L-citrulline<br>Methionine<br>4-aminobutanoate | Amino acid Degradation |
| Glycerol<br>Coenzyme A | Fatty acid and Lipid Degradation |

Metabolites were grouped based on their involvement in metabolic pathways.

## Discussion

Bacteria resist antibiotics by activating drug efflux pumps and/or enzymes which modify the antibiotic or its target [7]. Cellular metabolic rearrangements can also cause antibiotic resistance [46, 48-52]. In this report, we show that the antibiotic INH causes *Mtb* to increase the accumulation of cell surface, cellular, and extracellular polyP. Gallein, a bacterial PPK 1 and 2 inhibitor, inhibits *Mtb* cell surface, cellular, and extracellular polyP accumulation in both the presence and absence of INH, and in the presence of INH inhibits *Mtb* cell envelope formation in some but not all *Mtb* cells. Both INH and gallein have modest effects on metabolite levels, but the combination of INH and gallein strongly reduces levels of metabolites in several metabolic pathways. Possibly as a consequence of the effects of gallein on polyP levels, cell envelope formation, and metabolites, gallein inhibits *Mtb* growth in both *in vitro* culture and in human macrophages, and strongly potentiates INH inhibition of *Mtb* growth in *in vitro* culture and in human macrophages (Figure 5).

**Figure 5:**
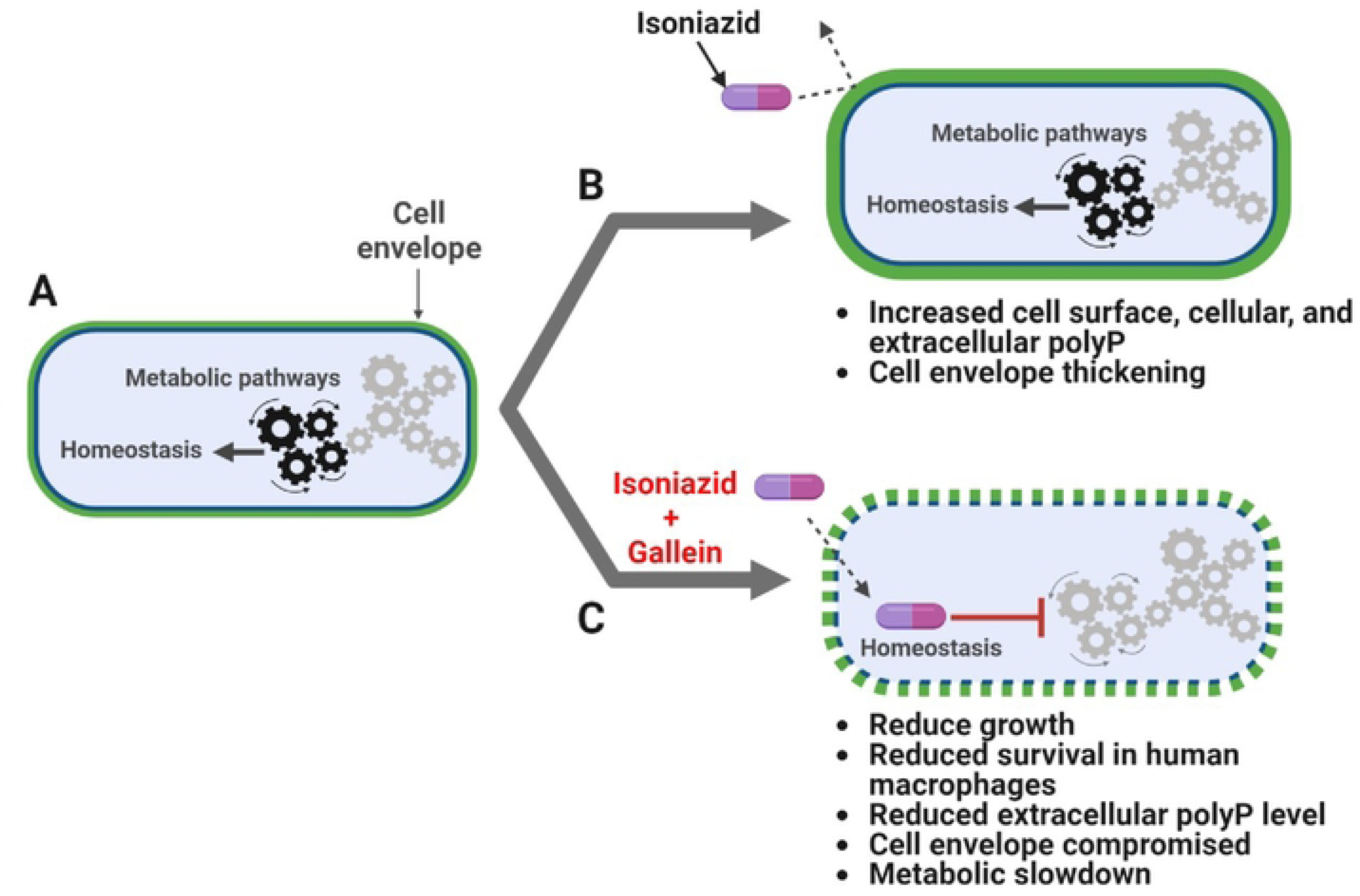
INH and gallein synergistically alter cellular homeostasis. A) *Mtb* under no antibiotic pressure maintain homeostasis. B) Under isoniazid (INH) pressure, *Mtb* grow slowly in *in vitro* culture, and increase the accumulation of cell surface, cellular, and extracellular polyP, as well as thickness of the cell envelope. C) When exposed to both INH and gallein, *Mtb* exhibit decreased metabolic activity, decreased cell surface and extracellular polyP, and decreased cell envelope formation and integrity in many cells, resulting in very slow growth in both *in vitro* culture and within macrophages. The pathway diagram was created using BioRender.com.

We observed a significant increase in cellular, extracellular, and cell surface polyP in response to INH. Compared to log-phase cells, stationary phase *Mtb* have more polyP and are more resistant to killing by INH [27, 53], and capsular polyP accumulation protects *N. gonorrhoeae* from antibiotics [23, 24]. Combining these observations, one possibility is that *Mtb* increase polyP in response to INH as a protective measure. The antibiotic rifampicin causes *Mtb* to thicken its capsular outer layer and increase the net negative charge of the cell surface to reduce rifampicin permeability [54]. It is possible that the INH-induced increase in the accumulation of the highly negatively charged polyP on the cell surface might similarly increase the net negative charge of the cell surface, reducing permeability to INH.

At 5 µM, gallein inhibits extracellular polyP levels in *Mtb* and inhibits INH-induced increases in cell surface and extracellular polyP. Gallein treatment of *P. aeruginosa* required concentrations exceeding 25 µM to reduce intracellular polyP accumulation, and 100 µM to mimic the effects of PPK deletion [33]. Our findings suggest that gallein exhibits a greater effectiveness on *Mtb* compared to *P. aeruginosa*.

*Mtb* can differentiate into heterogeneous populations, such as non-replicating persisters and growing bacteria with the capacity to become persisters [55]. *Mtb* treated with INH causes a rapid killing of cells followed by a reduction in the killing rate as the number of persister cells increases [56, 57]. In *Clostridiodes difficile,* a gram-positive bacteria, some cells in a population stochastically become antibiotic tolerant persisters [58]. *C. difficile* forms metabolically dormant spores that resist antibiotic pressure and persist in the host [59]. Two distinct morphotypes of *C. difficile* spores exist, one with thick outer surface layer and the other with thin outer surface layer [60, 61]. Multi-drug resistant *Mtb* have thicker cell envelopes [62]. INH caused many, but not all, *Mtb* cells to have thickened cell envelopes. In response to INH and gallein, *Mtb* formed two distinct populations of cells: one with cell envelope thicknesses comparable to INH-treated cells, and the other with no discernible cell envelope. One possibility is that under combined INH and gallein pressure, either stochastic differentiation or asymmetric cell division gives rise to ∼43% of cells with an intact cell envelope that have the potential to become slow growing persisters, and ∼ 57% of cells with compromised cell envelope that eventually die, and this might be the reason we observed no net increase in *Mtb* cell growth in the presence of 50 µM gallein and INH.

For control *Mtb* cells, there was a large variation in cell surface polyP levels, and INH increased both the average cell surface polyP levels and cell envelope thickness. At day 14, although there was a clear bimodal distribution of cell envelope thicknesses in the presence of INH and gallein, there was no observable bimodal distribution of cell-surface polyP levels. This suggests that there is not a strict correlation between cell-surface polyP level and cell envelope thickness, and that the effect of combined INH and gallein on cell wall thicknesses may be due to effects on additional pathways in addition to cell surface polyP.

*Mtb* uses the glucose disaccharide trehalose as a core component of cell surface glycolipids important for virulence, and during the transition into a non-replicating persister phase, it metabolizes trehalose to generate ATP [63-65]. In *Mtb* exposed to the combination of INH and gallein, the significant reduction in levels of trehalose and components necessary for trehalose biosynthesis, such as UDP-α D-glucose [66] suggests that the combination of INH and gallein compromises trehalose-dependent cell-surface glycolipid, and thus cell envelope, formation as we observed under TEM, and by decreasing trehalose levels, reduces this source of energy to prevent persister formation.

Gallein significantly reduced the levels of biotin. *Mtb* relies on biotin synthesis for its survival during infection, and the disruption of the biotin biosynthesis pathway results in cell death rather than growth arrest [67]. This suggests that gallein may inhibit *Mtb* growth by promoting cell death rather than merely inhibiting growth. The combination of gallein and INH resulted in decreased levels of metabolites associated with several key metabolic pathways. It is plausible that the synergistic action of INH and gallein on *Mtb* growth is not solely due to the reduced integrity of the cell envelope, but also to the disruption of other metabolic pathways.

In conclusion, the combination of INH and gallein affects several aspects of *Mtb* physiology, and gallein thus potentiates INH antibiotic effects on *Mtb*. Because they are not present in mammalian cells [31], PPKs are attractive target for suppressing *Mtb* growth and elimination.

## Materials and Methods

### Cell culture

Human peripheral blood was collected from healthy volunteers who gave written consent, and with specific approval from the Texas A&M University human subjects institutional review board. Peripheral blood mononuclear cells (PBMCs) were purified as previously described [68]. The PBMCs were cultured in RBCSG (RPMI-1640, # 15-040-CV, Corning, Corning, NY) containing 10% bovine calf serum (VWR Life Science Seradigm, Radnor, PA) and 2 mM L-glutamine (Lonza)), and where indicated containing 25 ng/mL human granulocyte-macrophage colony-stimulating factor (GM-CSF) ((# 572903, Biolegend, San Diego, CA) at 37 °C in a humidified chamber with 5% CO_2_ in 96-well, tissue-culture-treated, polystyrene plates (type 353072, Corning) with 2 x 10^5^ cells in 200 µL in each well. At day 7, loosely adhered cells were removed by gentle pipetting and removing the medium, and fresh RBCSG containing GM-CSF (as described above) was added to the cells to a final volume of 200 µL per well, and *Mtb* survival assays were performed as described below.

The attenuated (mc-ΔleuDΔpanCD) Biosafety Level-2 strain of *Mtb*, which is a derivative of the H37Rv strain [69], was a gift from Dr. Jim Sacchettini, Texas A&M University, College Station, TX and is referred to as *Mtb* in this paper. This strain was cultured according to the methods described in [70-72] in Middlebrook 7H9 broth (Becton, Dickinson and Company (BD), Sparks, MD) containing 0.5% glycerol (VWR), 0.05% Tween 80 (MP Biomedicals, Solon, OH), Middlebrook Oleic ADC Enrichment (BD), 50 μg/mL leucine (VWR), and 50 μg/mL pantothenate (Beantown Chemical, Hudson, NH) either in a rotator (10 rpm) or on plates with 7H10 agar (BD) and the above additives. All cultures were incubated at 37 °C in a humidified incubator. These supplemented cultures are hereafter referred to as 7H9S or 7H10S.

### *Mtb* growth assay

*Mtb* was grown in 24-well plates (type 353047, Corning). Each well was filled with 1 ml of 7H9S media, either containing no isoniazid (INH) or supplemented with 0.1, 1, 10, or 100 µg/ml INH (prepared from a 50 mg/ml stock in water, Cat#I3377, Sigma, Livonia, MI). *Mtb* samples obtained from log-phase liquid culture, as described above, were washed twice with 10 ml of 7H9S by centrifugation at 4000 x g for 10 minutes. The optical density at 600 nm (OD_600_) was measured in a well of a 96-well, tissue-culture-treated, polystyrene plate (type 353072, Corning) at 600 nM with a Synergy Mx monochromator microplate reader (BioTek, Winooski, VT). The *Mtb* were then resuspended in an appropriate volume of 7H9S to achieve a final OD_600_ of 1. Ten microliters of *Mtb* with an OD_600_ of 1 were added to each well to reach a final OD_600_ of 0.01 in 1 ml. The plates were subsequently incubated in a container with humidity provided by wet paper towels at 37 °C in a humidified incubator. On the 21st day, the *Mtb* culture was gently resuspended, and 100 µl of the cells were transferred to a 96-well, tissue culture-treated plate (# 353072, Corning). The OD_600_ was measured using a microplate reader. Given that we observed complete inhibition of *Mtb* growth with 100 µg/ml INH in our experimental setup, we opted to utilize 1 µg/ml INH for all our assays. This choice aligns with previous reports from other research groups, which had documented the resistance of *Mtb* to concentrations of INH greater than 1 µg/ml [73, 74].

To investigate the impact of gallein (3’,4’,5’,6’-Tetrahydroxyspiro[isobenzofuran-1(3H),9’- (9H)xanthen]-3-one) (Cat#3090, Tocris, Minneapolis, MN) and/or INH on *Mtb* growth, a *Mtb* culture plate was prepared as described method. However, in this case, each well of a type 353046, 6 well, tissue culture-treated plate (Corning) contained a final OD_600_ of 0.01 in 5 ml. *Mtb* was incubated with gallein at concentrations of 0.005, 0.05, 0.5, 5, or 50 µM and/or 1 µg/ml INH. A 50 mM gallein stock in DMSO (VWR) was diluted to 5 mM in 7H9S and further serially diluted in 7H9S to obtain lower concentrations. The control well contained 7H9S with DMSO, which was similarly serially diluted in 7H9S, as was done for gallein. The OD_600_ of the cells was measured daily for 14 days, and the *Mtb* growth curves were generated as a percentage of the OD_600_ on day 0.

To determine the viability of *Mtb* cells pre-exposed to INH and/or gallein for 14 days, 200 µl of cells from above experiments were washed twice with 1 ml of 7H9S, resuspended in 200 µl of 7H9S without INH and gallein 96-well plates ((# 353072, Corning), and the OD_600_ of the cells was measured daily for 14 days, and the *Mtb* growth curves were generated as a percentage of the OD_600_ on day 0.

### Bacterial survival assay

To determine the effect of INH and/or Gallein on the survival of *Mtb* in human macrophages, human macrophages (from blood monocytes cultured with GM-CSF for 6 days were mixed with *Mtb* following [27], in the absence or in the presence of 1 µg/ml INH and/or 5 µM Gallein. At day 7, after isolating monocytes from donor blood, after removing loosely adhered cells as described above, 200 µL RBCSGLP (RBCSG containing 50 μg/mL leucine and 50 μg/mL pantothenate) were added to macrophages in each well in 96-well, tissue-culture-treated, polystyrene plates (# 353072, Corning) and incubated for 30 minutes at 37 °C. Meanwhile, 1 mL of *Mtb* from a log phase culture was washed twice with RBCSGLP without GM-CSF by centrifugation at 12,000 x g for 2 minutes in a microcentrifuge tube, resuspended in 1 mL of RBCSGLP, and the OD_600_ of 100 µl of the culture in a well in a 96-well plates (# 353072 Corning) was measured as above. 200 µl of RBCSGLP was used as a blank. The bacteria were diluted to an OD_600_ of 0.5 (∼10^7^ *Mtb*/ mL) in RBCSGLP. *Mtb* (∼1 µl) was added to macrophages in each well such that there were ∼5 bacteria per macrophage, considering ∼20% of the blood monocytes converted to macrophages in the presence of GM-CSF [75]. The bacteria-macrophage co-culture plate was centrifuged at 500 x g for 3 minutes with a Multifuge X1R Refrigerated Centrifuge (Thermo Scientific, Waltham, MA) to synchronize phagocytosis of the bacteria, and incubated for 2 hours at 37 °C. The supernatant medium was removed by gentle pipetting and was discarded. 200 µL of PBS warmed to 37 °C was added to the co-culture in each well, cells were gently washed to remove un-ingested extracellular bacteria, the PBS was removed, and 200 µL of RBCSGLP with GMCSF in the absence or in the presence of 1 µg/ml INH and/or 5 µM gallein was added to the cells. After 2 hours, cells were washed twice with PBS as above. 200 µL of RBCSGLP with GMCSF in the absence or in the presence of 1 µg/ml INH and/or 5 µM gallein was then added to the cells. After 4 and/or 48 hours of infection, macrophages were washed as above with PBS, the PBS was removed, and cells were lysed using 100 µL 0.1% Triton X-100 (Alfa Aesar, Ward Hill, MA) in PBS for 5 minutes at room temperature by gentle pipetting, and 10 µl and 100 µL of the lysates were plated onto agar plates (as described above for *Mtb* culture). The *Mtb* containing agar plates were incubated for 3 to 4 weeks or until the *Mtb* colonies appeared. Bacterial colonies obtained from plating 10 µl and 100 µl lysates were manually counted, the number of viable ingested bacterial colonies per 10 µl and 100 µl lysates was calculated, and the number of viable ingested bacteria colony forming units (cfu) per ml of lysate was then calculated, which corresponds to the number of viable ingested bacteria in ∼2 x 10^5^ macrophages. To calculate the percent of control, cfu/ml of the control was considered 100%.

### PolyP assays, RNA extraction, and quantitative reverse transcription PCR

Log phase *Mtb* cultures were prepared similarly to the growth assays in 6-well, tissue culture-treated plates (# 353046, Corning), with the exception that 5 µM gallein and/or 1 µg/ml INH were used. Plates were incubated in a container humidified with wet paper towels at 37 °C for 24 hours. After incubation, 100 µl of cells were transferred to a 96-well, black/clear, tissue-culture-treated, glass-bottom plate (# 353219, Corning) for imaging as described below. The remaining cells were transferred to a 15-ml conical tube, harvested by centrifugation at 4000 x g for 10 minutes, and 4.5 ml of the supernatant was transferred to a new 15-ml conical tube for extracellular polyP measurement. Cell pellets were further processed for RNA extraction, following the procedure outlined below. PolyP levels in the supernatant were assessed by adding 25 µg/ml of DAPI (Biolegend) (from a stock of 2 mg/ml) and measuring fluorescence at 415 nm excitation and 550 nm emission, as previously described [76]. PolyP standards (Sodium Polyphosphates, Glassy, Spectrum, New Brunswick, NJ) at concentrations of 0, 0.5, 1, 10, 100, 200, and 500 µg/ml were prepared in 7H9S. The polyP content was normalized to the total protein content, which was determined from the cell lysates as described below.

For RNA extraction, cell pellets obtained from the treatments described above were resuspended in 250 μL of GITC lysis buffer (containing 4 M guanidine isothiocyanate and 50 mM Tris-HCl at pH 7). The cells were lysed by incubating at 95 °C for 10 minutes. Ten microliters of the lysates were used to determine the total protein content using a Bradford assay. The remaining lysates were used to extract RNA using an RNA extraction kit (Zymo Research, Irvine, CA). Complementary DNA (cDNA) was synthesized from 2 µg of RNA using the Maxima H Minus First Strand cDNA Synthesis kit (Thermo Scientific). Quantitative PCR was performed using SYBR GreenER™ qPCR SuperMix Universal reagent (Thermo Scientific), following the manufacturer’s instructions using a QuantStudio (TM) 6 Flex thermal cycler (Thermo Scientific). The levels of *Mtb’s ppk1* and *ppk2* mRNAs were determined using the gene-specific primers listed in Table 3.

### Fluorescence microscopy

To determine the localization of polyP in *Mtb*, 100 µl of *Mtb* cells from the growth assay on Day 21 or the growth assay on Day 1, 5, or 14 were transferred to a 96-well, black/clear, tissue-culture-treated, glass-bottom plate (# 353219, Corning) or cells smears were prepared on Superfrost micro glass slides (Cat#48311-703, VWR) as described previously [77]. The cells were fixed with 4% (wt/vol) paraformaldehyde (Cat#19210, Electron Microscopy Sciences, Hatfield, PA) in PBS for 10 minutes. After fixation, the cells were washed two times with 300 µl of PBS, blocked with 1 mg/ml BSA (Thermo Scientific) in PBS, and then stained with 10 µg/ml of GFP-PPX (provided generously by Dr. Ursula Jacob from the University of Michigan) in PBS/0.1% Tween 20 (PBST; Fisher Scientific) [42]. Following staining, the *Mtb* cells were washed three times with PBST, and 200 µl of PBS was added to the well, or coverslips were mounted on slides with smears with Vectashield hard set mounting medium (Cat# H-1800, Vector Labs, Burlingame, CA) and left to dry overnight in darkness. Images of *Mtb* were captured using a 100× oil-immersion objective on a Nikon Eclipse Ti2 (Nikon, Kyoto, Japan), and image deconvolution was performed using the Richardson–Lucy algorithm [78] in NIS-Elements AR software (Nikon). The integrated fluorescence density was measured in randomly selected individual cells manually using the freehand selection feature in Fiji (ImageJ) [79].

### Transmission electron microscopy

*Mtb* cells treated with 5 µM gallein and/or 1 µg/ml INH for 14 days were prepared as described for the growth assays above. A volume of 100 µl of cells in 7H9S was fixed by adding an equal volume of 2× fixative, which contained 84 mM NaH_2_PO_4_, 68 mM NaOH, 4% paraformaldehyde (Cat#19210, Electron Microscopy Sciences), and 1% glutaraldehyde (Cat#0875, VWR). The samples were gently rocked for 1 hour and then stored at 4°C. Sample preparation for TEM imaging was performed by the Texas A&M University Microscopy and Imaging Center Core Facility’s staff (RRID: SCR_022128). Briefly, on the following day, the fixed samples were collected by centrifugation for 5 minutes at 14,000 × g. Subsequently, they were postfixed and stained for 2 hours with 1% osmium tetroxide in 0.05 M HEPES at pH 7.4. The samples were then collected by centrifugation and washed with water five times, and dehydrated with acetone according to the following protocol: 15 minutes in 30%, 50%, 70%, and 90% acetone each, followed by three changes of 100% acetone, each lasting 30 minutes. During the final wash step, a minimal amount of acetone was retained, just enough to cover the pellets, to prevent rehydration of the samples. Subsequently, the samples were infiltrated with modified Spurr’s resin (Quetol ERL 4221 resin; Electron Microscopy Sciences; RT 14300) in a Pelco Biowave processor (Ted Pella, Inc., Redding, CA). The process included 1:1 acetone-resin for 10 minutes at 200 W (no vacuum), 1:1 acetone-resin for 5 minutes at 200 W (vacuum at 20 inches Hg, with vacuum cycles involving open sample container caps), and 1:2 acetone-resin for 5 minutes at 200 W (vacuum at 20 inches Hg). This was followed by four cycles of 100% resin for 5 minutes each at 200 W (vacuum at 20 inches Hg). The resin was then removed, and the sample fragments were transferred to BEEM conical-tip capsules that were prefilled with a small amount of fresh resin. More resin was added to fill the capsules, and they were left to stand upright for 30 minutes to ensure that the samples sank to the bottom. The samples were polymerized at 65°C for 48 hours in an oven and then left at room temperature (RT) for an additional 24 hours before sectioning. Sections of 70 to 80 nm thickness were obtained using a Leica UC/FC7 ultramicrotome (Leica Microsystems), deposited onto 300-mesh copper grids, and stained with uranyl acetate-lead citrate. Grids were imaged using a JEOL 1200 EX TEM operating at 100 kV. Cell wall thickness was measured using ImageJ.

### Metabolic activity assay

Cell proliferation and viability in *Mtb* can be assessed by incubating the cells with resazurin reagent for 6-12 hours, and monitoring the color change from blue to purple [80]. Metabolically active cells transform the non-fluorescent blue dye (resazurin) into a fluorescent pink product (resorufin), while inactive cells rapidly lose their metabolic capacity and, consequently, do not generate a fluorescent signal [43]. *Mtb* from the log phase culture were prepared as previously described for the proliferation assay, with 100 µl of cells being prepared per well in 96-well, tissue culture-treated plate (# 353072, Corning). *Mtb* was treated with 5 µM gallein and/or 1 µg/ml INH, and the plate containing the *Mtb* was incubated at 37 °C for 24 hours. Cells were then incubated with prewarmed Deep Blue Cell Viability resazurin dye (Biolegend) to a final concentration of 10% in each well for 12 hours [80]. The fluorescence signal was then measured using a microplate reader following the manufacturer’s protocol.

### Metabolomics

*Mtb* cells from log-phase cultures were prepared as described for the growth assays, with the exception that 10 ml of culture was prepared. The *Mtb* cells were treated with 5 µM gallein and/or 1 µg/ml INH. After 24 hours, 10 ml of the culture was collected by centrifugation at 4000 x g for 10 minutes at 4 °C. The cells were then washed twice with 10 ml of chilled (0–4°C) phosphate-buffered saline (PBS) to prevent metabolite contamination from the culture media. The cells were resuspended in 10 ml of PBS [81], and 100 µl of this suspension was transferred to a 96-well plate to measure the OD_600_. The remaining cell suspension was centrifuged again at 4000 x g for 10 minutes at 4 °C to collect the cell pellets for metabolite extraction. After the final wash, excess PBS was carefully removed from all the samples. The day before the assay, 10 ml of extraction solvent (acetonitrile (BDH83639.400, VWR): methanol (BDH20864.400, VWR):water (40:40:20)) was prepared and stored at ™70°C. On the day of the experiment, heavy amino acid standards (Metabolomics amino acid mix standard, Cat# MSK-A2-1.2, Cambridge Isotope Laboratories, Tewksbury, MA) were added as a spike to the extraction solvent to 5 µM final concentration. To halt bacterial metabolism, the *Mtb* pellets were immediately suspended in 200 µl of spiked extraction solvent that had been pre-cooled on dry ice [46]. *Mtb* disruption was achieved using a Mini-beadbeater-16 (BioSpec Products, Bartlesville, OK). The *Mtb* samples (200 µl) were placed in 2 ml Polypropylene Microvials (Cat#10832, BioSpec Products) containing approximately 70 µl of 0.1 mm Zirconia/Silica beads (Cat# 11079101z, BioSpec Products). Bead-beating was performed in three cycles of 1 minute at 3,450 oscillations/min, with 2 minutes of cooling on ice between cycles [82]. Subsequently, the microvials containing the disrupted cells and zirconium beads were incubated at ™20°C for 20 minutes. They were then clarified by centrifugation at 8,000 x g for 15 minutes at 4 degrees Celsius. The resulting supernatant was filtered through a 0.22 µm filter (Cat# UFC30GV25, Merck Millipore, Cork, IRL) into 0.2 ml glass Stepvial inserts (Cat# 200 238, ThermoScientific). These step vials were placed inside 0.25 ml polypropylene vials with polypropylene caps with PTFE/silicone septa (Cat# 200 410, ThermoScientific). The pellets containing beads were resuspended in 100 µl of Radioimmunoprecipitation assay buffer (RIPA) (Cat#89900, Thermo Scientific) containing 1X protease and phosphatase inhibitor cocktail (Cat#1861281, Thermo Scientific), incubated on ice for 15 minutes, clarified by centrifugation at 10,000 x g for 5 minutes at 4°C, and 25 µl of the supernatant was used to determine the protein amount following the manufacturer’s instructions using a Pierce BCA Protein Assay Kit (Cat#23225, Thermo Scientific). A vial containing a quality control mixture pool of 12.5 µl from each sample (150 µl total) was prepared along with the other samples. These vials were sealed with Parafilm and stored at ™80 °C before the untargeted metabolomics analysis [81]. Samples were analyzed with a Shimadzu high-performance liquid chromatography (HPLC) (Nexera X2 LC-30AD, Kyoto, Japan) coupled to a Sciex TripleTOF 6600 high-resolution mass spectrometer (HRMS) for the separation and detection of various classes of metabolites in the samples using an untargeted metabolomics approach at the UTSW metabolomics core facility (https://www.utsouthwestern.edu/research/core-facilities/metabolomics/). Metabolites detected in at least three biological replicates were considered for further analysis. The peak area of metabolites was normalized to the total protein content determined from *Mtb* lysates, and pathway enrichment analysis was conducted using the online analytical tool Metaboanalyst (www.metaboanalyst.ca) and the BioCyc Database, employing the *Mtb* H37Rv reference genome [83].

### Statistical analysis

Statistical analyses were performed using Prism 10 (GraphPad Software, Boston, MA) or Microsoft Excel. P < 0.05 was considered significant.

## Contact for Reagent and Resource Sharing

Further information and requests for reagents may be directed to, and will be fulfilled by, the authors Ramesh Rijal or Richard Gomer.

## Acknowledgments

This work was supported by National Institutes of Health grant GM139468.

## Author Contributions

R. R. designed and performed experiments, analyzed data, and wrote the paper, and R. H. G. coordinated the study, wrote the paper, and acquired funding.

## Supporting information

**Figure S1:** INH reduces *Mtb* growth in *in vitro* culture. (A - C) *Mtb* cultures were grown for 14 days in the absence (Control) or presence of 1 µg/ml INH and/or 0.005 µM (A), 0.05 µM (B), or 0.5 µM (C) gallein. The OD_600_ was measured daily, and growth was determined as a percentage of Day 0 OD_600_. (D) *Mtb* were cultured for 21 days in the presence of the indicated concentrations of INH, and the OD_600_ was measured on Day 21. (E) *Mtb* at day 14 from Figures 1A and 1B were washed, regrown in the absence of INH or gallein for 14 days, and the OD_600_ was measured daily. Growth was determined as a percentage of the day 0 OD_600_. All values are mean ± SEM of three independent experiments. * P < 0.05; ** P < 0.01; *** P < 0.001 (One-way ANOVA with Dunnett’s multiple comparisons test).

**Figure S2:** INH and gallein increase percentages of *Mtb* cells with reduced cell surface polyP and cellular debris accumulation. (A) The percentages of cells with cell surface polyP levels less than 20 fluorescence units from Figure 2J. All values are mean ± SEM of three independent experiments. * P < 0.05 (One-way ANOVA with Tukey’s multiple comparisons test, and t-tests between Control and the combination of INH and Gallein. (B) Differential interference contrast (DIC) images of *Mtb* from Figure 2J. Representative images from at least three independent experiments are shown. Bars are 10 µm for main images and 5 µm for inset. White arrows indicate cellular debris.

**Figure S3:** INH or gallein does not alter *ppk1* and *ppk2* mRNA levels. Total RNA was extracted from *Mtb* cultures grown for 24 hours in the absence or presence of 1 µg/ml isoniazid (INH) and/or 5 µM gallein. The RNA was reverse-transcribed to generate cDNA, and the levels of *ppk1* and *ppk2* cDNA were quantified by quantitative PCR using gene-specific primers (Supplemental Table 1). The cDNA level from untreated *Mtb* (Control) was set to 1. All values are mean ± SEM of three independent experiments.

**Figure S4:** Gallein inhibits *Mtb* metabolic activity. (A) *Mtb*, treated without or with 1 µg/ml INH and/or 5 µM gallein for 24 hours, were incubated with cell viability dye for 12 hours, and fluorescence was measured. The average of the control was considered 100%. (B) The score plot of the two principal components (2 and 3) of a principal component analysis model, built on the entire metabolomics dataset (Figure 4) from three independent experiments, is color-coded by group with confidence ellipses. Control datasets are red circles, INH-treated datasets are purple circles, gallein-treated datasets are green circles, and INH and gallein-treated datasets are blue circles. The contribution ratios (variance) of the two principal components are shown in parentheses. All values represent the mean ± SEM of three independent experiments for (A). **** P < 0.001 (One-way ANOVA with Tukey’s multiple comparisons test).

**Table S1:** Oligonucleotides for quantitative real-time polymerase chain reaction (qPCR).

